# Susceptibility to diet induced obesity at thermoneutral conditions is independent of UCP1

**DOI:** 10.1101/2021.06.30.450595

**Authors:** Sebastian Dieckmann, Akim Strohmeyer, Monja Willershäuser, Stefanie Maurer, Wolfgang Wurst, Susan Marschall, Martin Hrabe de Angelis, Ralf Kühn, Anna Worthmann, Marceline M Fuh, Joerg Heeren, Nikolai Köhler, Josch K. Pauling, Martin Klingenspor

**Affiliations:** Chair for Molecular Nutritional Medicine, Technical University of Munich, TUM School of Life Sciences, Freising, Germany; EKFZ - Else Kröner-Fresenius Center for Nutritional Medicine, Technical University of Munich, Freising, Germany; ZIEL - Institute for Food & Health, Technical University of Munich, Freising, Germany; Institute of Developmental Genetics, Helmholtz Zentrum München, Germany; Technische Universität München-Weihenstephan 85764 Neuherberg/Munich, Germany; German Center for Neurodegenerative Diseases (DZNE), Site Munich, Germany; Institute of Experimental Genetics, Helmholtz Zentrum München, Germany; Chair of Experimental Genetics, TUM School of Life Sciences, Technical University of Munich, Freising, Germany; German Center for Diabetes Research (DZD), Neuherberg, Germany; Institute of Developmental Genetics, Helmholtz Zentrum München, Germany. Max-Delbrück-Center for Molecular Medicine in the Helmholtz Association, Berlin, Germany; Department of Biochemistry and Molecular Cell Biology, University Medical Center Hamburg-Eppendorf, Hamburg, Germany; LipiTUM, Chair of Experimental Bioinformatics, TUM School of Life Sciences, Technical University of Munich, Freising, Germany

## Abstract

**Objective:** Activation of uncoupling protein 1 (UCP1) in brown adipose tissue (BAT) upon cold stimulation leads to substantial increase in energy expenditure to defend body temperature. Increases in energy expenditure after a high caloric food intake, termed diet-induced thermogenesis, are also attributed to BAT. These properties render BAT a potential target to combat diet-induced obesity. However, studies investigating the role of UCP1 to protect against diet-induced obesity are controversial and rely on the phenotyping of a single constitutive UCP1-knockout model.

To address this issue, we generated a novel UCP1-knockout model by Cre-mediated deletion of Exon 2 in the UCP1 gene. We studied the effect of constitutive UCP1 knockout on metabolism and the development of diet-induced obesity.

**Methods:** UCP1 knockout and wildtype mice were housed at 30°C and fed a control diet for 4-weeks followed by 8-weeks of high-fat diet. Body weight and food intake were monitored continuously over the course of the study and indirect calorimetry was used to determine energy expenditure during both feeding periods.

**Results:** Based on Western blot analysis, thermal imaging and noradrenaline test, we confirmed the lack of functional UCP1 in knockout mice. However, body weight gain, food intake and energy expenditure were not affected by deletion of UCP1 gene function during both feeding periods.

**Conclusion:** Conclusively, we show that UCP1 does not protect against diet-induced obesity at thermoneutrality. Further we introduce a novel UCP1-KO mouse enabling the generation of conditional UCP1-knockout mice to scrutinize the contribution of UCP1 to energy metabolism in different cell types or life stages.

## 1 Introduction

Thermogenic brown adipose tissue (BAT) is the main contributor to non-shivering thermogenesis, the process to maintain normothermia in a variety of small mammals. Non-shivering thermogenesis is mediated by the uncoupling protein 1 (UCP1), which enables high rates of oxygen consumption by the mitochondrial electron transport chain without ATP production. The most potent stimulation of UCP1 and non-shivering thermogenesis is mediated by the sympathetic innervation of BAT with the neurotransmitter norepinephrine triggering beta-3-adrenergic receptor signaling in brown adipocytes. Furthermore, meal-associated thermogenesis in BAT is activated upon food intake (Glick, Teague, and Bray 1981) by the prandial surge of the gut peptide hormone secretin (Li et al. 2018). Other activators of BAT thermogenesis have been reported (Zietak and Kozak 2016; Gnad et al. 2014) and recent findings suggest that brown fat conveys effects on systemic metabolism and energy balance by means of paracrine intercellular and endocrine interorgan crosstalk. The beneficial metabolic effects of BAT clearly go beyond the combustion of calories (Kajimura, Spiegelman, and Seale 2015). Activation of meal-associated thermogenesis in BAT initiates meal termination and thereby contributes the control of energy intake (Schnabl, Li, and Klingenspor 2020). Together with the potential of BAT to impact energy balance and systemic metabolism by clearing glucose from circulation, these characteristics render UCP1 and BAT potential targets to improve cardiometabolic health and the treatment of type 2 diabetes (Becher et al. 2021).

In this context the question whether UCP1 can protect against diet induced obesity (DIO) has been studied repeatedly. Standard housing temperature (20-23°C) represents a mild cold challenge for laboratory mice resulting in a two-fold increase of daily energy expenditure (Fischer, Cannon, and Nedergaard 2018). BAT is the source for this thermoregulatory heat production. UCP1 knockout mice when kept at standard housing temperature do not develop diet-induced obesity (Bond and Ntambi 2018) and even seem to be protected, having lower body weight than WT mice (Liu et al. 2003; T. Wang et al. 2008; Keipert et al. 2020). It has been proposed that this is due to alternative thermogenic mechanisms less efficient than UCP1-dependent thermogenesis (Liu et al. 2003; T. Wang et al. 2008; Keipert et al. 2020), that are recruited to cope with the need for thermoregulatory heat production in UCP1-KO mice.

Housing mice at higher temperatures (27°C – 30°C), corresponding to their thermoneutral zone eliminates this heat sink. Since UCP1 in BAT is inactive at thermoneutrality, the lack of thermogenic BAT function should be without consequences for energy balance. Indeed, several studies using the established UCP1-KO mouse model originally generated by Leslie Kozak and coworkers confirmed this expectation (Enerbäck et al. 1997; Liu et al. 2003; Zietak and Kozak 2016; Winn et al. 2017; Maurer et al. 2020; Fischer et al. 2020). In contrast, other studies reported that UCP1 knockout mice are more susceptible to diet induced obesity at thermoneutrality (Feldmann et al. 2009; von Essen et al. 2017; Rowland et al. 2016; Luijten et al. 2019; Pahlavani et al. 2019). One explanation for the increased susceptibility to DIO in UCP1-KO mice may be adaptations in metabolism, leading to a more efficient metabolism or the lack of diet-induced thermogenesis in UCP1-KO mice (von Essen et al. 2017). However, recent data from a UCP1 knockdown model (H. Wang et al. 2021) demonstrate that UCP1 abundance alone does not protect against DIO at thermoneutrality. Despite having remarkable reduced but still activatable UCP1 levels, these mice are not more or less prone to DIO compared to wildtype littermates with normal functional levels of UCP1.

This showcases the urgent need for new UCP1-KO models to scrutinize the role of UCP1 on energy balance and metabolism. So far two UCP1 knockout models are available (Enerbäck et al. 1997; Bond and Ntambi 2018). Other transgenic mice with impaired UCP1 expression include knockdown models (Chen, Hsu, and Huang 2018; H. Wang et al. 2019) or diphtheria toxin chain A induced depletion of UCP1 expressing cells (Lowell et al. 1993; Rosenwald et al. 2013).

In the present study we therefore introduce and validate a novel Cre-mediated UCP1-KO model and demonstrate that deletion of UCP1 has no effect on energy balance regulation at thermoneutrality.

## 2 Material and Methods

### 2.1 Animal model

The UCP1 knockout mouse line was generated in frame of the EUCOMM program and is a constitutive UCP1 knockout model on a C57BL/6N background (Skarnes et al. 2011; Pettitt et al. 2009). It originates from the UCP1^tm1a^ mouse, carrying a lacZ & neomycin cassette, two FLP sites and three loxP sites (Figure 1 A). Through crossing with a FLP mouse, the lacZ and neomycin cassette as well as one FLP site and one loxP site are removed, resulting in a UCP1^tm1c^ (UCP1-WT) mouse. Cross breeding this mouse with a Rosa26-CRE mouse results in the UCP1^tm1d^ (UCP1-KO) mouse carrying a germline deletion of the exon 2 of the UCP1 gene. UCP1-KO and UCP1-WT mice were crossed to generate UCP1-HET mice. The UCP1 knockout line, is maintained by crossing male and female UCP1-HET mice. All studied mice were derived of our heterozygous maintenance breeding. Mice were bred and housed at 23°C ambient temperature with a 12/12 h light/dark cycle and had libitum access to water and chow diet.

**Figure 1:**
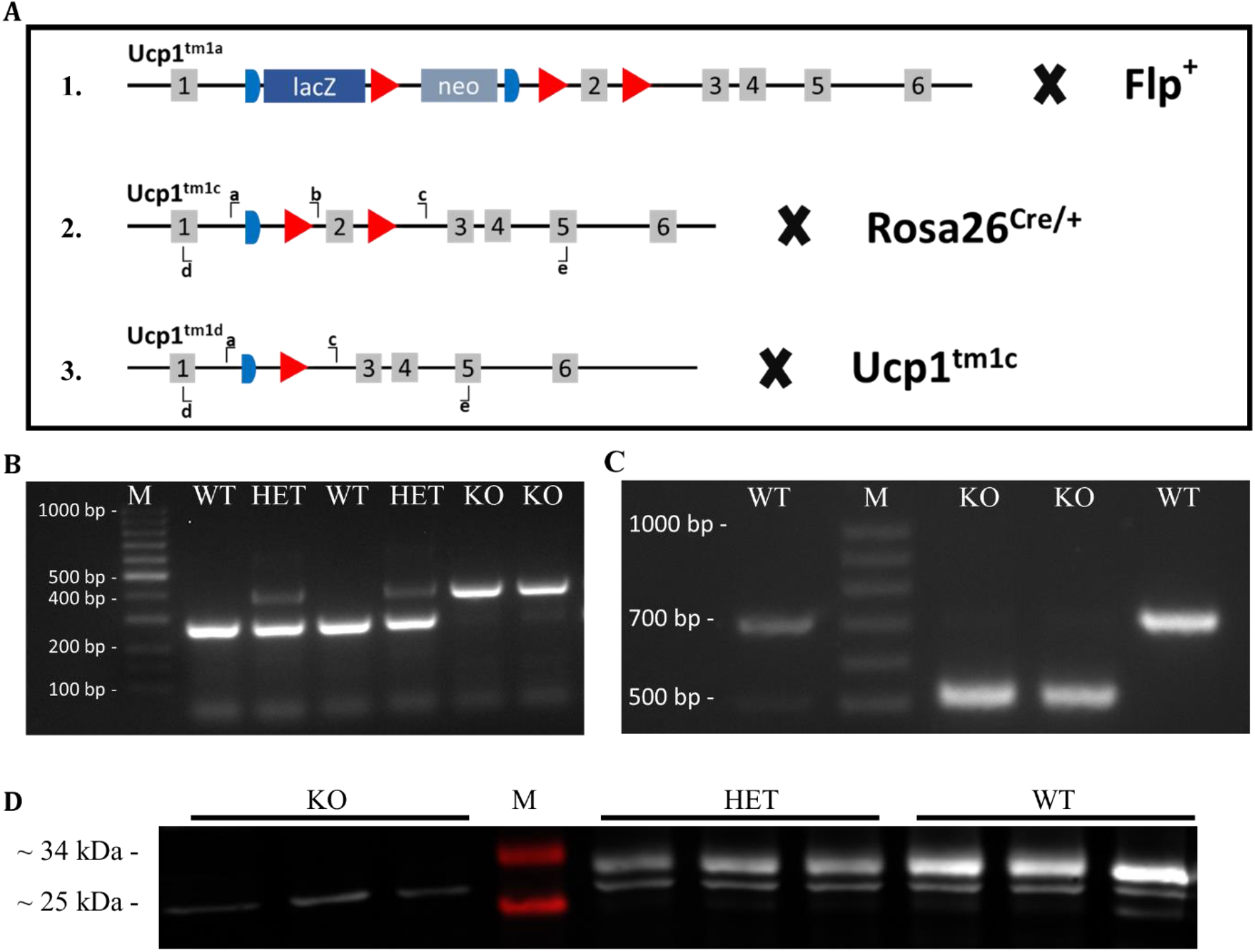
Overview and validation of the Ucp1 knockout strategy. (**A**) Breeding scheme for the generation of the Ucp1 knockout. (**A1**) Ucp1tm1a mice containing three loxP sites (red), two frt sites (blue) as well as a lacZ and a neo cassette were crossed with mice expressing flippase (Flp+). (**A2**) The resulting Ucp1tm1c (WT) mouse is crossed with a Rosa26CRE/+ mouse, deleting exon 2 of the ucp1 gene, generating Ucp1tm1d (KO) mice. (**A3**) Ucp1tm1c and Ucp1tm1d mice are crossed to generate HET mice. Lower case letters indicating binding positions of primers used for PCR (a-c) and RT-PCR (d&e). (**B**) PCR of gDNA from tissue samples of WT, HET and KO mice. (**C**) RT-PCR products from iBAT of WT and KO mice. (**D**) Representative western blot analysis for Ucp1 (~33 kDA) in KO, HET and WT mice. See Supplementary Figure 2 for the uncropped image.

All animal experiments were performed according to the German animal welfare law and approved by the district government of Upper Bavaria (Regierung von Oberbayern, reference number ROB-55.2-2532.Vet_02-15-128).

#### 2.1.1 HFD feeding at thermoneutrality

Male wildtype (n = 7) and knockout (n = 7) mice for the high fat diet feeding experiment were obtained from our heterozygous maintenance breeding. At the age of 8 weeks, mice were switched from chow to a chemically defined control diet with a fat content of 50 g/kg (CD, ~13 kJ% from Fat, 15.3 MJ/kg, Snifff Cat. No S5745-E702). Simultaneously, mice were single caged and transferred to climate cabinets with an ambient temperature of 30 °C and 55 % RH. After an acclimatization phase of 4 weeks, mice were switched from CD to a high fat diet with a fat content of 250 g/kg (HFD, ~48 kJ% from fat, 19.6 MJ/kg, Snifff Cat. No S5745-E712). After 8 weeks of HFD feeding, mice were killed by CO_2_ asphyxiation. Whole blood was taken by cardiac puncture, collected in lithium heparin-coated tubes (Sarstedt, Nümbrecht/Germany), and centrifuged at 4°C for 10 min with 1500 x g. The plasma supernatant was transferred to fresh tubes and snap frozen in liquid nitrogen. Subsequently, cecal content and tissues were dissected, weighed, and immediately snap frozen in liquid nitrogen. Cecal content, tissues and plasma were stored at −80°C until further processing. Body weight and food intake were determined twice a week between 12.00 PM and 4.00 PM. Additionally, body composition was determined every other week by nuclear magnetic resonance spectroscopy (mq7.5, Bruker BioSpin GmbH, Rheinstetten/Germany). Mice were maintained on a 12/12 h light/dark cycle and had ad libitum access to water and the respective diets during the whole experiment. Food was replaced completely twice a week to avoid rancidity of the HFD at 30°C. Energy expenditure, energy intake, energy excretion and metabolic efficiency of CD and HFD-fed mice were assessed as described below.

### 2.2 Indirect calorimetry, basal metabolic rate and noradrenaline tests

Indirect calorimetry was performed based on an open respirometer system (LabMaster System; TSE Systems, Bad Homburg/Germany) similar to previously described methodology (Maurer et al. 2015). O_2_ consumption and CO_2_ production were determined after 2.5 weeks of feeding CD and after 4 weeks of HFD. Mice were transferred in specially equipped cages in a climate cabinet (KPK 600, Feutron, Germany) set to 30°C after determining body weight and body composition in the afternoon (2.00 – 5.00 PM). The measurement was started on the next day at 6.00 AM (CD) or 12.00 PM (HFD) and continued two (CD) or three (HFD) dark phases. The air from the cages was extracted over a period of 1 min every 4-6 min. Heat production was calculated according to (Heldmaier 1975) as: *HP*[*mW*] = (4.44 + 1.43 *∗ respiratory exchange ratio*) *∗ oxygen consumption* [*ml*/*h*]

Basal metabolic rate (BMR) was determined immediately after the last night phase of the indirect calorimetry measurement of the HFD period. Mice were deprived of food between 7.00 – 8.00 AM for at least 4 hours. BMR was calculated as the mean of the four lowest consecutive heat production measurements during the last 90 minutes of fasting, similar to a previous published method (Fromme et al. 2019). Subsequently, noradrenaline tests were performed between 10.00 AM – 5.00 PM at 26°C to avoid noradrenaline induced hypothermia. Noradrenaline (1 mg/kg, Arterenol®, Sanofi) was injected intraperitoneally. Air was extracted continuously from the cages with a measurement period of 1 min over 60 min.

### 2.3 Collection of food spillage and faeces

Embedding material was collected form cages after indirect calorimetry for each mouse separately, to correct food intake for spillage and to determine energy loss by faecal excretion. Material was dried at room temperature under a chemical flow hood for at least 1 week. Subsequently, cage material was fractionated based on size by shaking the material on a sieve shaker (EML 200 Digital Plus T, Haver & Boecker, Oelde/ Germany) for 5 min with an interval of 0.5 min at an amplitude of 1.4, through sieves with different mesh sizes (4, 3.15, 2.5, 1.25 and 1 mm, VWR International GmbH, Darmstadt/Germany). Flowthrough of the 1 mm sieve was collected in a pan. Each sieve was scanned for spilled food and faeces (the majority of faeces will be present in the 1.25 mm sieve). If applicable, food and faeces were picked with tweezers and collected for weighing and determination of energy content by bomb calorimetry. Food intake (in grams) during the indirect calorimetry sessions was corrected for the amount of collected food spillage (in grams).

### 2.4 Determination of energy content of food and faeces by bomb calorimetry

The energy content of the diets and the faecal pellets collected during indirect calorimetry was determined with an isoperibolic bomb calorimeter (Model Nr. 6400, Parr Instrument Company, IL/USA). Energy content of the diets was determined on food samples collected at different time points during the experiment (CD n = 9, HFD n = 10). Energy intake was calculated by multiplying the mean energy content of the diets (kJ/g) with the amount of food intake (in grams).

The collected faeces was weighed and grinded with metal balls for 2.5 min at 30 Hz (Tissue Lyser II, Retsch GmbH. Haan/Germany). Grinded faeces was pressed into a pellet, weighed, and subjected to bomb calorimetry. Benzoeic acid (~0.7 g) was added as combustion aid. Energy lost via faeces was calculated for each mouse by multiplying the total amount of faeces collected (in grams, see 2.4) by the energy content (kJ/g) determined by bomb calorimetry.

### 2.5 Thermal imaging

Thermal imaging was performed as described previously (Maurer et al. 2015) with 1-3 day old new-born pups. In brief, at least 3 serial pictures were taken of each litter in 6-well cell culture plates (T890 thermal imager, Testo, Lenzkirch/Germany). Image analysis was performed with the IRSoft Software (version 4.6, Testo, Lenzkirch/Germany) and the temperature above the interscapular BAT deport (interscapular skin surface temperature, iSST) was determined.

### 2.6 Genotyping

Genotyping was performed on earpieces obtained during tagging of the animals. Tissues were lysed (10 mM TRIS, 50 mM KCl, 0.45 % Nonidet P40, 0.45 % Tween-20, 10 % gelatin in H20 at pH 8.3% with 0.2 mg/ml Proteinase K) for 4 h at 65°C and vigorous shaking. Proteinase K was inactivated by heating for 10 min at 95°C. PCR (denaturation: 5 min / 95°C followed by 39 amplification cycles with 30s / 95°C, 45s / 54°C, 45s / 72°C and a final elongation 10 min / 72°C) was performed with three primers (Figure 1 A, “a”: AAGGCGCATAACGATACCAC, “b”: TACAATGCAGGCTCCAAACAC, “c”: CGAGCACAGGAAGTTCAACA, Eurofins Genomics, Ebersberg/Germany) and the ImmoMix*™* kit (Bioline, Cat. No BIO-25020) according to the manufacturer’s instructions.

### 2.7 RNA isolation and cDNA synthesis and sequencing

RNA precipitation was performed with TRIsure*™* (Bioline, London/UK) following to the manufacturer’s instructions, from deep-frozen iBAT. Precipitated RNA was loaded to spin columns (SV Total RNA Isolation System, Promega, Cat# Z3105), centrifuged for 1 min with 12,000 x g and further processed according to the supplier’s instructions. RNA concentration was determined spectrophotometrically (Infinite 200 PRO NanoQuant, Tecan). cDNA synthesis was performed with 1 µg RNA (SensiFAST*™* cDNA Synthesis Kit, Bioline, Cat# BIO-65053), according to the manufacturer’s instructions. PCR (denaturation: 10 min / 95°C followed by 30 amplification cycles with 1 min / 95°C, 30s / 54 °C, 40s / 72°C and a final elongation 5 min / 72°C) was performed with primers (“d”: cggagtttcagcttgcctggca, “e”: tcgcacagcttggtacgcttgg, Eurofins Genomics, Ebersberg/Germany, Figure 1 A) and products were separated by gel electrophoresis on a 1 % agarose gel. Separated PCR products were visualized under a UV light, cut, immediately weighed and stored at −20°C. PCR products were purified with the Wizard SV Genomic DNA Purification System (Promega, Cat# A2361), and sent in to a commercial sequencing platform (Eurofins Genomics, Ebersberg/Germany). Analysis of sequencing results was performed with the “Benchling” platform (https://www.benchling.com/).

### 2.8 Protein expression analysis by SDS-Page and Western Blot

Protein was isolated from interscapular BAT, homogenized in 10 µl/mg isolation buffer (50 mM Tris, 1% NP-40, 0.25% sodium deoxycholate, 150 mM NaCl, 1 mM EDTA) containing 0.1 % phosphatase (Sigma-Aldrich, St. Louis MO/USA) and 0.1 % protease inhibitor cocktail (Sigma-Aldrich, St. Louis MO/USA) with a dispersing device (Miccra D-1, Miccra GmbH, Heitersheim/Germany). The homogenized samples were centrifuged 15 min at 4°C with 14.000 rcf. The clear layer of the supernatant was isolated by pipetting and centrifuged again. Samples were cleared from residual fat by a second extraction of the clear phase with a syringe. Protein concentrations were determined with the Pierce*™* BCA Protein Assay Kit (ThermoScientific, Rockford IL/USA) according to the manufacturer’s instructions. For protein detection, 30 µg protein were separated in a 12.5 % SDS-PAGE and transferred to a nitrocellulose membrane. Subsequently, primary antibody was applied to detect UCP1 (ab23841, Abcam, UK, 1:5000) followed by primary antibody detection using an IR-dye conjugated secondary antibody (IRDye 800CW, LI-COR, Lincoln NE/USA, 1:20000). The IR signal was detected with the Azure Sapphire*™* biomolecular imager (azure biosystems, Dublin CA/USA). Image analysis was conducted with the Image Studio*™* Lite software version 5.2.

### 2.9 DNA Extraction and 16S rRNA Sequencing

Cecal contents were collected together with other tissues and immediately snap frozen in liquid nitrogen and stored at −80°C. DNA isolation, library preparation and sequencing were performed at the ZIEL – Core Facility Microbiome of the Technical University of Munich. Briefly, DNA was extracted using previously published protocols (Klindworth et al. 2013). For the assessment of bacterial communities primers specifically targeting the V3-V4 region of the bacterial 16S rRNA (Forward-Primer (341F-CCTACGGGNGGCWGCAG; Reverse-Primer (785r-ovh): GACTACHVGGGTATCTAATCC) gene including a forward and reverse illumina specific overhang and a barcode were used. Sequencing was performed using an Illumina MISeq DNA platform. Obtained multiplexed sequencing files have been analyzed using the IMNGS platform, which is based on the UPARSE approach for sequence quality check, chimera filtering and cluster formation (Edgar 2013; Lagkouvardos et al. 2016). For the analysis standard values for barcode mismatches, trimming, expected errors and abundance cutoff have been used and only sequences between 300 and 600 bp were considered for analysis. Downstream analysis of the IMNGS platform output files were performed using the RHEA R pipeline (Lagkouvardos et al. 2017). In brief, obtained abundances have been normalized and quality of obtained sequences was assessed using rarefaction curves (McMurdie and Holmes 2014). Analysis of alpha diversity, beta diversity and group comparisons have been performed using default settings. Exceptions have been applied for group comparisons for zOTUs and taxonomic levels (abundance cutoff 0.5 and exclusion of alpha diversity measures). Graphical output was modified for presentation using inkscape (https://inkscape.org). Assignment of zOTUs to taxons has been performed using the SILVA database (Version 138.1 (Quast et al. 2013)). Assignment of species to specific zOTUs with EZBioCloud (Yoon et al. 2017).

### 2.10 Lipid Extraction and Mass Spectrometry Analysis

Lipid extraction for quantitative analysis using Lipidyzer*™* platform (SCIEX) was done using an adapted Methyl-tert-butyl-ether (MTBE) extraction protocol. Lipidyzer*™* internal standards mixture was prepared according to the manufacturer’s instruction but dissolved in MTBE. To each 50 µL plasma aliquot, 50 µL water; 50 µL Internal Standard, 500 µL MTBE and 160 µL Methanol was added, shortly vortexed and incubated on a mixer for 30 minutes. 200 µL of water was added and centrifuged at 16000g. The supernatant was transferred in vials and the residual phase re-extracted using MTBE: Methanol: Water in the ratio 3:1:1. The collected supernatants were evaporated with a vacuum centrifuge and resuspended in 250 µL of 10 mM ammonium acetate in Dichloromethane: Methanol (50:50 (v/v)).

Samples were analyzed using a QTRAP 5500 (AB SCIEX) equipped with Differential Mobility Spectrometer (DMS) interface (Schneider et al. 2010) operating with SelexION technology, coupled to a Shimadzu Nexera X2 liquid chromatography system. The Lipidyzer platform*™* was operated via the software Analyst version 1.6.8 and Lipidomics workflow manager (SCIEX). A detailed description of this shotgun approach has been previously reported (Lintonen et al. 2014). The Lipidyzer*™* Platform was tuned using the SelexION Tuning Kit (SCIEX) according to the manufacturer’s recommendations and a system suitability test was performed using the System Suitability Kit (SCIEX) according to the manufacturer’s instructions. The Lipidyzer*™* Platform uses 10 mM ammonium acetate in Dichloromethane: Methanol (50:50 (v/v)) as running buffer, Dichloromethane: Methanol (50:50 (v/v)) as rinse 0&1, 2-propanol as rinses 2&3, and 1-propanol as a DMS modifier. 50µl of samples were injected for each of the two MRM methods: One with a DMS on and one with DMS off. MRM data acquisition, processing, and quantification was performed automatically by the lipidyzer lipidomics workflow manager. Lipid concentrations are given in nmol/ml.

### 2.11 Data analysis and statistics

General data analysis was performed with R (version 4.0.3) within R-Studio (version 1.3.1093). Unless otherwise indicated data are represented as means ± sd or with single values for each mouse. Student’s t-tests were performed with the package “ggpubr” (version 0.4.0). Anova, and linear model analysis with the package “stats” (version 4.0.3). Trapezoid area under the curves were calculated using the AUC function of the package “DescTools” (version 0.99.38).

Analysis of alpha diversity was performed with Prism 6 (GraphPad Software Inc., La Jolla CA/USA) using non-parametric Mann-Whitney U test. Beta diversity is visualized using non-metric multi-dimensional scaling based on generalized UniFrac and tested for significance using PERMANOVA. Differences in zOTUs have been determined using Kruskal-Wallis rank sum test with adjustments for multiple testing using the Benjamini & Hochberg method.

Multi-omics analysis was performed using R (version 4.0.4) and python (version 3.8.5). Multi-omics factor analysis (MOFA) (Argelaguet et al. 2018; 2020) was used for unsupervised data integration of the lipidome and microbiome data. The mofapy2 python package (version 0.5.8) and the MOFA2 R package (version 1.0.1) (for downstream analysis) were used together with custom visualization tools. Data integration analysis for biomarker discovery using latent variable approaches for ‘omics studies (DIABLO) (Singh et al. 2019) was used as a supervised analysis framework. DIABLO generalizes (sparse) partial least-squares discriminant analysis (PLS-DA) for the integration of multiple datasets measured on the same samples. For DIABLO analyses the mixOmics R package (version 6.14.0) (Rohart et al. 2017) was used along with custom code for randomized performance estimation.

## 3 Results

### Deletion of UCP1 Exon 2 leads to a loss of protein expression

The original mouse used to generate the knockout of UCP1 in the mice used in this study was generated in frame of the EUCOMM program via a “knockout first allele” approach (Skarnes et al. 2011; Pettitt et al. 2009). In order to generate WT (*UCP1*^*tm1c*^), the lacZ and the neomycin resistance cassette were removed from the UCP1^*tm1a*^ allele, by cross breeding with flippase expressing mice (Figure 1 A1), thus generating WT (*UCP1*^*tm1c*^) mice. These mice containing only one flippase recognition target (frt) and two lox P sites flanking exon 2 of the UCP1 gene were crossed with mice expressing Cre-recombinase under the control of the rosa26 promotor (Rosa26Cre/+) (Figure 1 A2). This resulted in the generation of KO (*UCP1*^*tm1d*^) mice by constitutive germline deletion of exon 2 in the UCP1 gene (Figure 1 A3). Deletion of exon 2 was first confirmed by PCR on genomic DNA with one forward and two reverse primers binding to distinct sites of the UCP1 gene (Figure 1 A2 & A3). As predicted, this resulted in a short (263 bp) product for WT mice (Figure 1 B, primers a-b) and a longer (388 bp) product for KO (Figure 1 B, primers a-c), while HET mice showed both products. Of note, the 1255 bp product generated by the primers a and c in WT and HET is not seen, as the elongation period of the PCR protocol is too short to produce a product of this size. To further investigate the consequences of exon 2 deletion, we performed a RT-PCR on RNA isolated from brown adipose tissue of both WT and KO mice. For the primer pair binding in exon 1 (d) and exon 5 (e) of the UCP1 gene (Figure 1 A), KO showed a smaller product size (~500 bp) compared to WT mice (~700 bp), as predicted by in-silico PCR (KO: 508 bp, WT: 707 bp, https://genome.ucsc.edu/cgi-bin/hgPcr) (Figure 1 C). Subsequent sequencing of the WT and KO PCR-products revealed that the deletion of exon 2 causes a frame shift, leading to a premature stop codon in exon 3 (Supplementary Figure 1). Consequently, KO mice do not express UCP1 protein, as confirmed by western blot analysis (Figure 1 D, Supplementary Figure 2).

### Thermogenic deficiency leads to decreased body weight in young KO mice

The loss of the major protein responsible for non-shivering thermogenesis resulted in a clear reduction of interscapular skin surface temperature (iSST) in newborn KO compared to WT mice (Figure 2 A&B). The loss of one functional UCP1 allele (HET) on the other hand had no implication on iSST in newborn pups compared to WT mice (Figure 2 A&B). The genotype distribution of offspring from HET/HET breeding pairs (generation F2-F3) did not significantly deviate from the mendelian distribution of 1:2:1 (Figure 2 C, Table 1). However, KO mice had lower body weight at weaning (at the age of ~3-4 weeks) compared to HET and WT mice (Figure 2 D), a phenotype that could be confirmed in the conventional UCP1-KO mouse on 129S1/SvImJ-but not on C57Bl/6J-background (Supplementary Figure 3 A&C). Irrespectively of the knockout model bodyweight of all three genotypes were similar at ~ 8 weeks of age (Figure 2 E, Supplementary Figure 3 B&D).

**Table 1:**
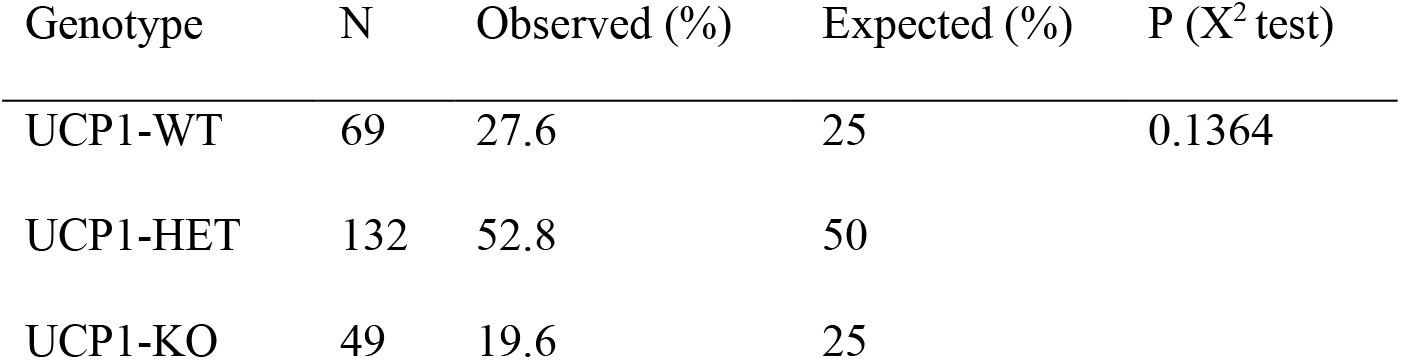
Offspring genotype distribution of heterozygous breeding pairs.

**Figure 2:**
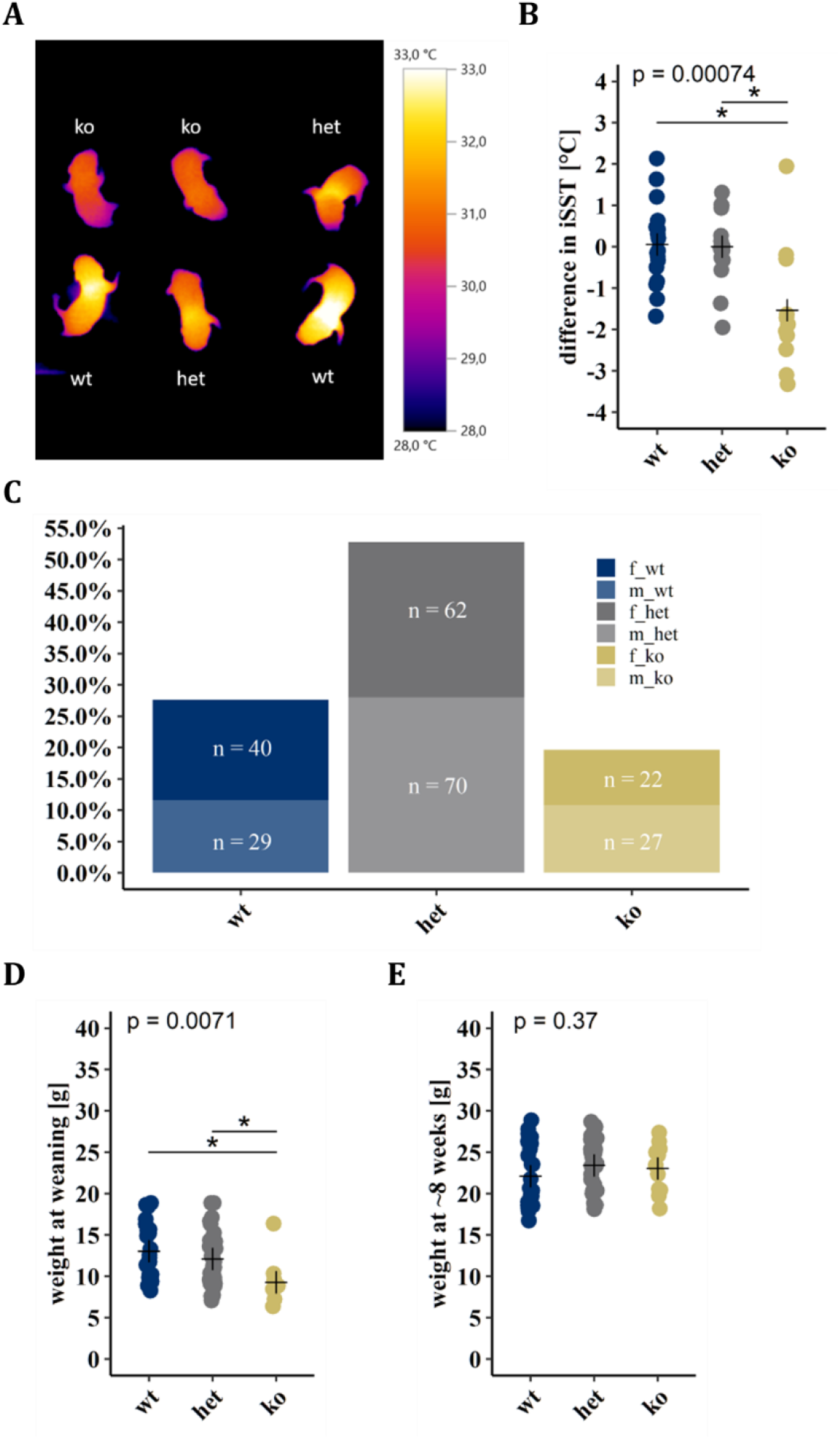
Lack of Ucp1 leads to phenotypic alterations in young mice. (**A**) Representative thermal image of newborn pups (2-3 days) of a Ucp1-HET breeding pair. (**B**) Analysis of interscapular skin surface temperature (iSST, n(wt) = 17, n(het) = 17, n(ko) = 11, N = 45 of 5 litters). (**C**) Offspring genotype distribution of Ucp1-HET breeding pairs (n = 15) at 23 °C ambient temperature (N = 250). (**D-E**) Body weight of female and male Ucp1-WT (n = 23), Ucp1-HET (n = 33) and Ucp1-KO (n = 11) mice (**D**) at weaning and (**E**) at the age of 8-weeks (N = 67 of 9 litters). (B,D,E) Crosses indicating group means. 1-Way ANOVA and t-test with bonferroni adjusted p-value, * = p-value < 0.05.

In summary this suggests a strain dependent effect of UCP1 depletion on early body weight that recovers with age.

### UCP1-KO and WT mice have similar susceptibility to DIO at thermoneutrality

The susceptibility UCP1-KO mice to diet induced obesity (DIO) under thermoneutral conditions is still a matter of debate. We addressed this controversial question using our novel UCP1-KO model by feeding mice at thermoneutrality a control diet (CD) for 4 weeks followed by 8 weeks of high-fat diet (HFD). At the start of the experiment at the age of 8 weeks, mice of both genotypes had similar body weights (data not shown). Cumulative body weight gain increased with time but was similar between both genotypes (Figure 3 A) as indicated by linear model analysis during CD (Duration P < 0.001, Genotype P = 0.491) and HFD (Duration P < 0.001, Genotype P = 0.188) feeding. In line, total energy intake between both genotypes was similar during control diet (CD) and high fat diet (HFD) feeding (Figure 3 B&C).

**Figure 3:**
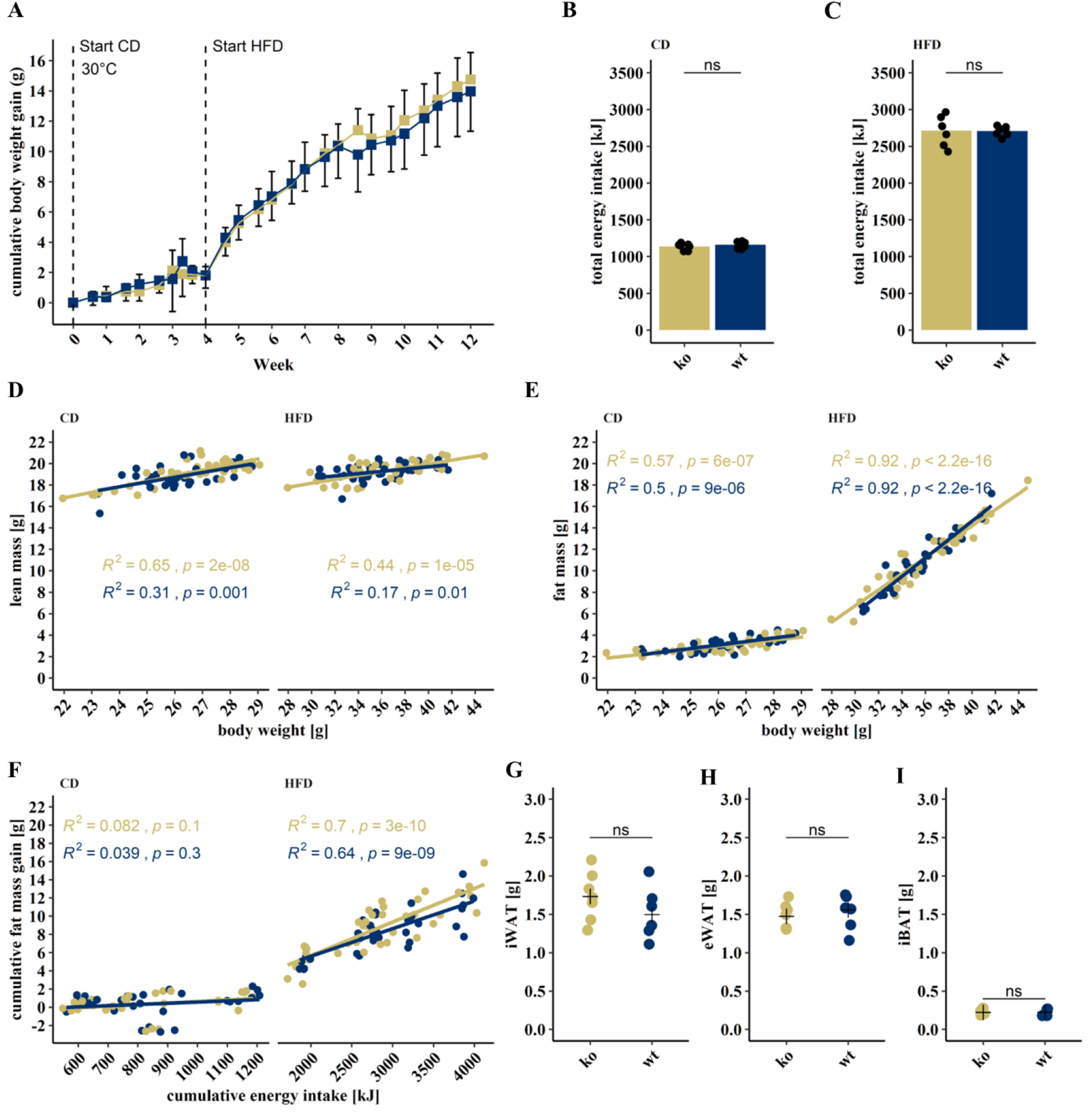
Similar susceptibility to diet induce obesity in UCP1-KO and WT mice. (**A**) Body weight of Ucp1-WT (wt, n = 7) and Ucp1-KO (ko, n = 7) mice at 30°C fed a control (CD) or high-fat diet (HFD). Total energy intake of mice during (**B**) CD and (**C**) HFD feeding. (D-E) Pearson correlation coefficient between measurements of (**D**) lean mass and body weight and (**E**) fat mass and body weight during CD (left) and HFD (right) feeding. (**F**) Metabolic efficiency in terms of correlation (Persons correlation coefficient) between cumulative fat mass gain and cumulative energy intake for CD (left panel) and HFD (right panel). Weights of dissected (**G**) inguinal white adipose tissue (iWAT), (**H**) epididymal white adipose tissue (eWAT) and (**I**) interscapular brown adipose tissue (iBAT) at the end of HFD feeding. (B,C,G-I) Student’s t-test ns = p-value > 0.05. Group means indicated as (B,C) bars and (G-I) crosses.

We determined body composition in terms of lean and fat mass at different time points of the experiment. Both lean mass (Figure 3 D) and fat mass (Figure 3 E) correlated well with body weight during both feeding regimes, with fat mass being the main contributor to the increase in body weight during HFD feeding (R^2^ > 0.9), in both WT and KO mice (Figure 3 D & E).

UCP1 knockout mice have been described to be metabolically more efficient (Feldmann et al. 2009; von Essen et al. 2017; Luijten et al. 2019), thus incorporating more fat mass per unit of energy intake. We addressed this question by linear model analysis of cumulative fat mass gain versus cumulative energy intake over the experimental period (Figure 3 F). There was no difference in the correlation of fat mass gain and energy intake between genotypes, consequently both UCP1-WT and UCP1-KO mice showed similar metabolic efficiency. Of note, this result was confirmed by determining metabolic efficacy as the percentage of food energy stored as fat mass, as described previously (von Essen et al. 2017) (Supplementary Figure 4 A&B). The similarity in fat mass of both UCP1-WT and UCP1-KO determined by NMR was reinforced by dissected weights of iWAT, eWAT and iBAT (Figure 3 G-I). Collectively these data analyses demonstrate that UCP1 ablation neither affected energy intake, nor body adiposity, nor metabolic efficacy when mice were kept at thermoneutral conditions.

### Plasma lipid composition of UCP1-KO and UCP1-WT mice is comparable

Activated BAT can clear substantial amounts of lipids from circulation (Heine et al. 2018; Bartelt et al. 2011). To study whether UCP1 ablation affected systemic lipid metabolism, a targeted lipidomic approach on plasma samples was performed. Lipid class composition was similar between both genotypes. Only cholesteryl esters (CE) were significantly more abundant in UCP1-WT compared to UCP1-KO mice (Figure 4 A). Concentration of CE, ceramides (CER), hexosylceramides (HCER) and sphingomyelins (SM) were significantly higher in UCP1-WT mice (Figure 4 B). However, fold changes (FC) between to UCP1-KO were rather small (CE, FC_wt/ko_ = 1.13; CER, FC_wt/ko_ = 1.17; HCER FC_wt/ko_ = 1.16; SM, FC_wt/ko_ = 1.1). The similarity of plasma lipid composition between both genotypes was confirmed by principal component analysis (PCA) of composition (Figure 4 C) and concentration (Figure 4 D) on a lipid species level.

**Figure 4:**
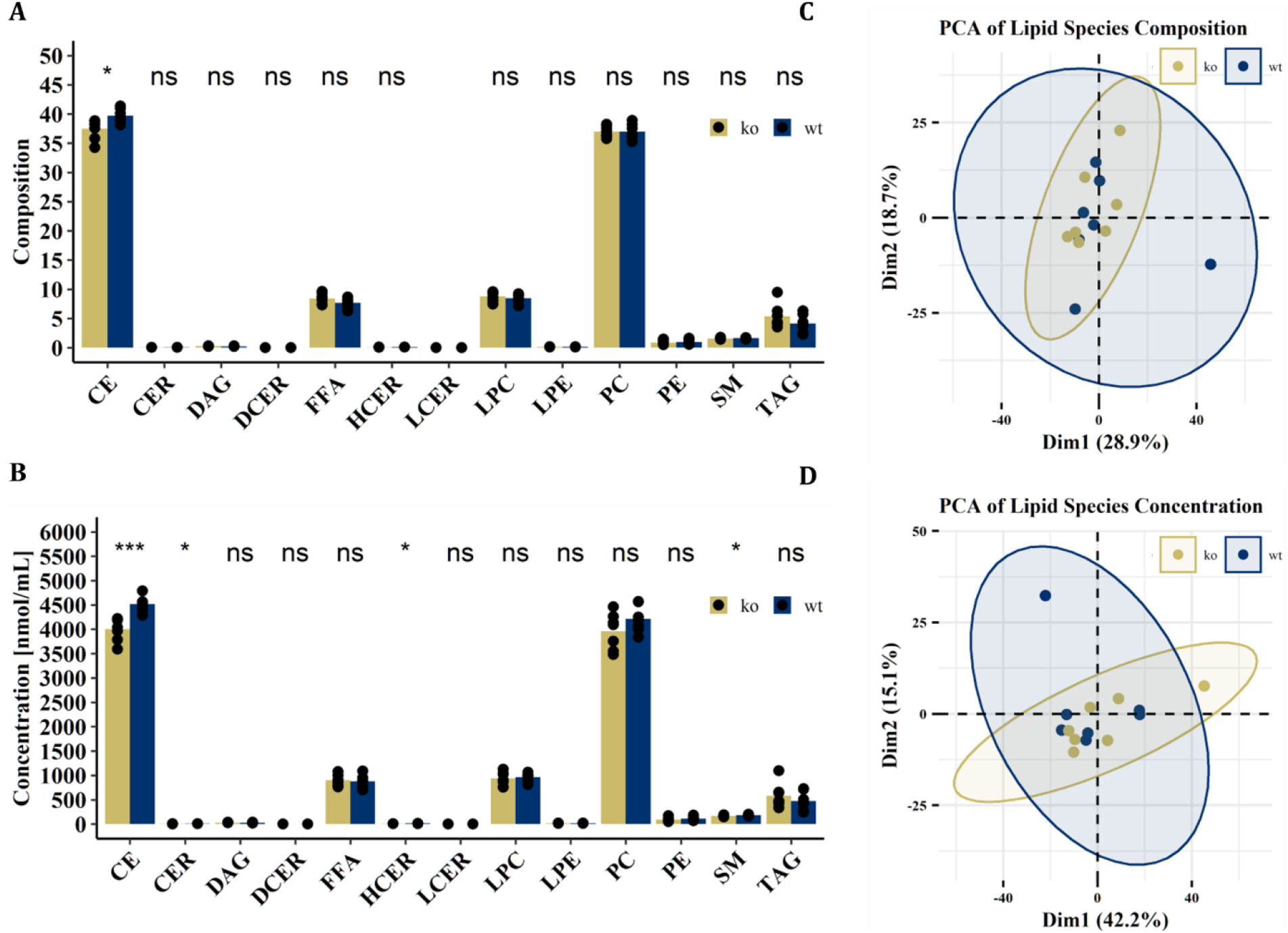
Plasma lipid profiles are comparable between UCP1-KO and UCP1-KO. **(A)** Composition and **(B)** concentration of lipid classes in plasma of UCP1-KO (ko, n =7) and UCP1-WT (wt, n = 7) mice housed at 30°C after 8 weeks of high fat diet feeding. Principal component analysis (PCA) of lipid species **(C)** composition and **(D)** concentration. Cholesteryl esters (CE), ceramides (CER), diacylglycerols (DAG), dihydroceramides (DCER), free fatty acids (FFA), hexosylceramides (HCER), lactosylceramides (LCER), lysophosphatidylcholines (LPC), lysophosphatidylethanolamines (LPE), phosphatidylcholines (PC), phosphatidylethanolamines (PE), sphingomyelins (SM). triacylglycerols (TAG). (A&B) Student’s t-test ns = p-value > 0.05, * = p-value < 0.05, *** = p-value < 0.001. Group means indicated as (A&B) bars.

Collectively, these data demonstrate that ablation of UCP1 did have only minor effects on steady state systemic lipid metabolism at thermoneutrality.

### Lack of UCP1 is associated with the abundance of specific microbial genera

The gut microbiome influences host metabolism (Tremaroli and Bäckhed 2012) and studies demonstrate an effect of microbiome composition on UCP1 expression (Wu et al. 2019) and thermogenesis (Ziętak et al. 2016). Consequently, we investigated whether UCP1 expression alters microbiome composition, by comparing the cecal microbiomes of UCP1-WT and UCP1-KO mice. Similar bacterial richness (alpha-diversity) was observed between genotypes by 16S rRNA analysis (Figure 5 A). However, deletion of UCP1 affected cecal microbial composition demonstrated by differences in beta-diversity between genotypes (Figure 5 B). Detailed analysis of the microbial composition revealed four zOTU significantly different between UCP1-KO and UCP1-WT based on unadjusted Kruskal-Wallis rank sum test (Figure 5 C-F). After adjustment for multiple comparisons, two of these zOTUs demonstrated a trend to higher abundance in UCP1-KO while the other were significantly more abundant in UCP1-WT mice. These zOTUs could be assigned to *Parabacteroides goldsteinii* (zOTU3 & zOTU4) and *Desuflovibrio fairfieldensis* (zOTU17 and zOTU19), respectively. Interestingly, *P. goldsteinii* has previously been reported to decrease in HFD induced obesity and diabetes and increase UCP1 expression in iBAT and iWAT in C57BL/6J mice (Wu et al. 2019). These data indicate a connection between UCP1 expression and the gut microbiota and confirm *P. goldsteinii* as a potential species associated with UCP1.

**Figure 5:**
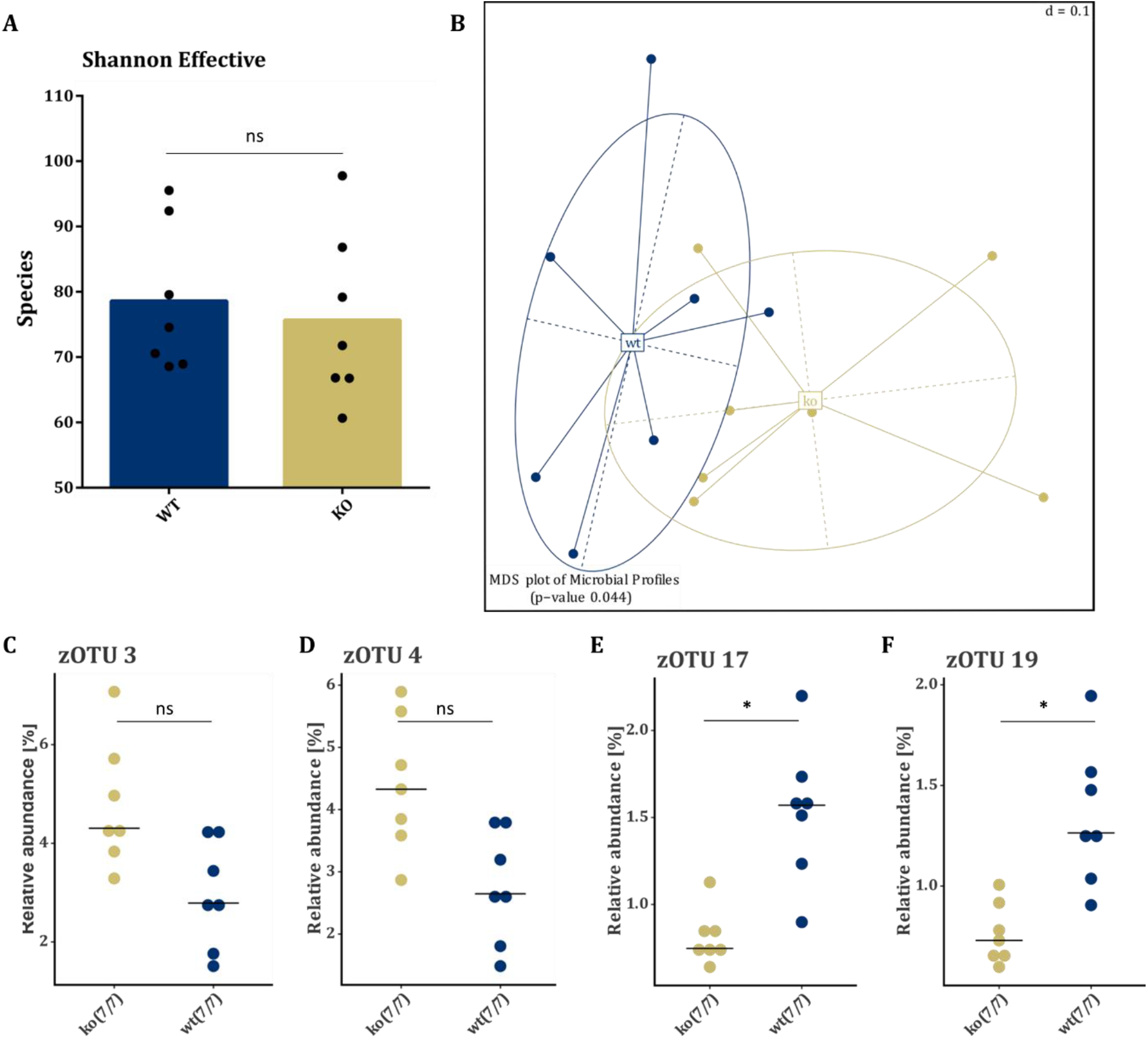
Single cecal microbial genera are associated with the presence of UCP1. Analysis of cecal microbiome of UCP1-KO (ko, n = 7) and UCP1-WT (wt, n = 7) mice housed at 30°C after 8 weeks of high fat diet feeding. Comparison of **(A)** alpha-diversity determined by Shannon effective index and **(B)** beta-diversity assessed by principal coordinates analysis. (**C-F)** Relative abundance of zOTU identified by statistically different unadjusted Kruskal-Wallis rank sum test between WT and KO mice. Statistical differences tested by (A) non-parametric Mann-Whitney U test ns = p-value > 0.05, (B) Permutational multivariate analysis of variance, (C-F) Kruskal-Wallis rank sum test with the Benjamini & Hochberg adjustment ns = p-value > 0.05, * = p-value < 0.05, Group means indicated as (A) bars, (C-F) lines.

The microbiome can substantially influence host lipid metabolism (Schoeler and Caesar 2019). As we identified small changes in both lipid metabolism and microbiome composition, we investigated potential interactions between microbiome and lipidome. Therefore, we analyzed the combined lipidome and microbiome data set using supervised (DIABLO PLS-DA) and unsupervised (MOFA) approaches. DIABLO PLS-DA revealed two sets of features that discriminated between UCP1-KO and UCP1-WT mice (Figure 6 A). However, quality assessment by repeated analysis of the dataset with randomly assigned groups (1000 iterations) demonstrated similar good discrimination as between UCP1-KO and UCP1-WT mice (Figure 6 A). Consequently, it was not possible to discriminate the observed difference between UCP1-KO and UCP1-WT from random differences between samples. This assumption was confirmed by unsupervised MOFA demonstrating no separation of the two genotypes by the two factor groups explaining the highest proportion of variance between UCP1-KO and UCP1-WT mice (Figure 6 B).

**Figure 6:**
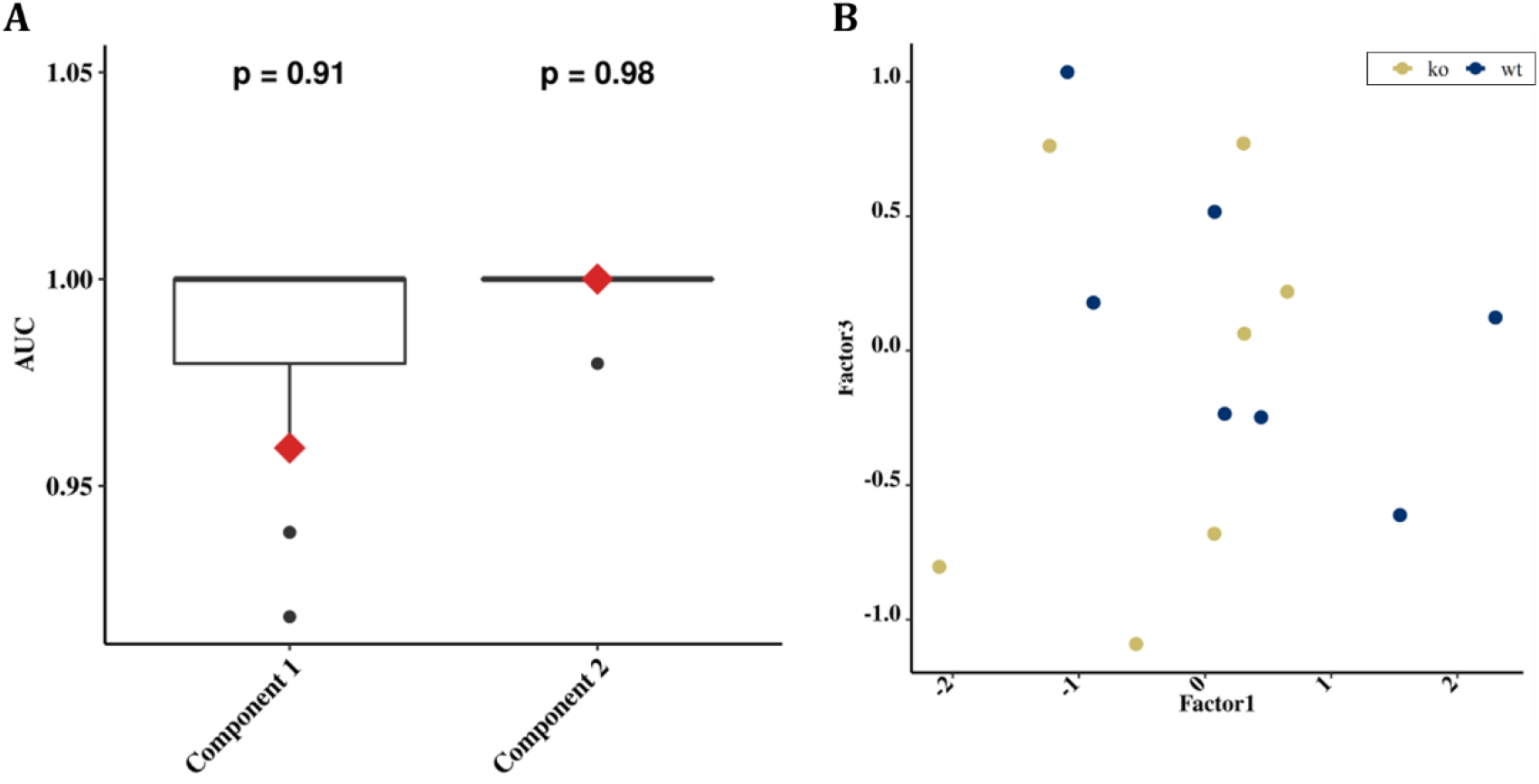
Multi-omics reveal no interaction between microbiome and lipidome explaining differences between UCP1-KO and UCP1-WT mice. Integrated analysis of the combined lipidome and microbiome data sets. **(A)** Receiver operating characteristic area under the curve (AUC) of data integration analysis for biomarker discovery using a latent components partial least squares discriminant analysis (DIABLO PLS-DA). Red diamonds indicate results of supervised DIABLO PLSA-DA of the components 1 and 2 (Comp1, Comp2). Box plots indicate results of 1000 randomized DIALBO PLS-DA analyses. P-value = #(AUC_sup_ < AUC_ran_) / 1000. **(B)** Visualization of the two factors explaining most of the variance between UCP1-KO and UCP1-WT mice based on multi-omics factor analysis (MOFA).

Consequently, no genotype specific interactions between plasma lipid composition and the microbiome were identified.

### UCP1-KO mice have similar energy balance at thermoneutrality

The effect of UCP1 knockout on energy balance regulation was investigated in detail by indirect calorimetry measurements 3-4 weeks after the start of CD or HFD feeding. We observed a clear diurnal pattern of the respiratory exchange ratio during CD feeding, being higher during the dark phase compared to the light phase (Figure 7 A&B) indicating that mice utilized more carbohydrates during the nocturnal activity phase while relying more on fatty acid metabolism during the daytime resting phase. During HFD feeding the respiratory exchange ratio was generally reduced compared to the CD period, demonstrating a shift in substrate utilization towards fatty acid oxidation based in the high fat content of the diet (Figure 7 C&D). However, no differences in respiratory exchange ratio between KO and WT mice were detected during either feeding period (Figure 7 B&D).

**Figure 7:**
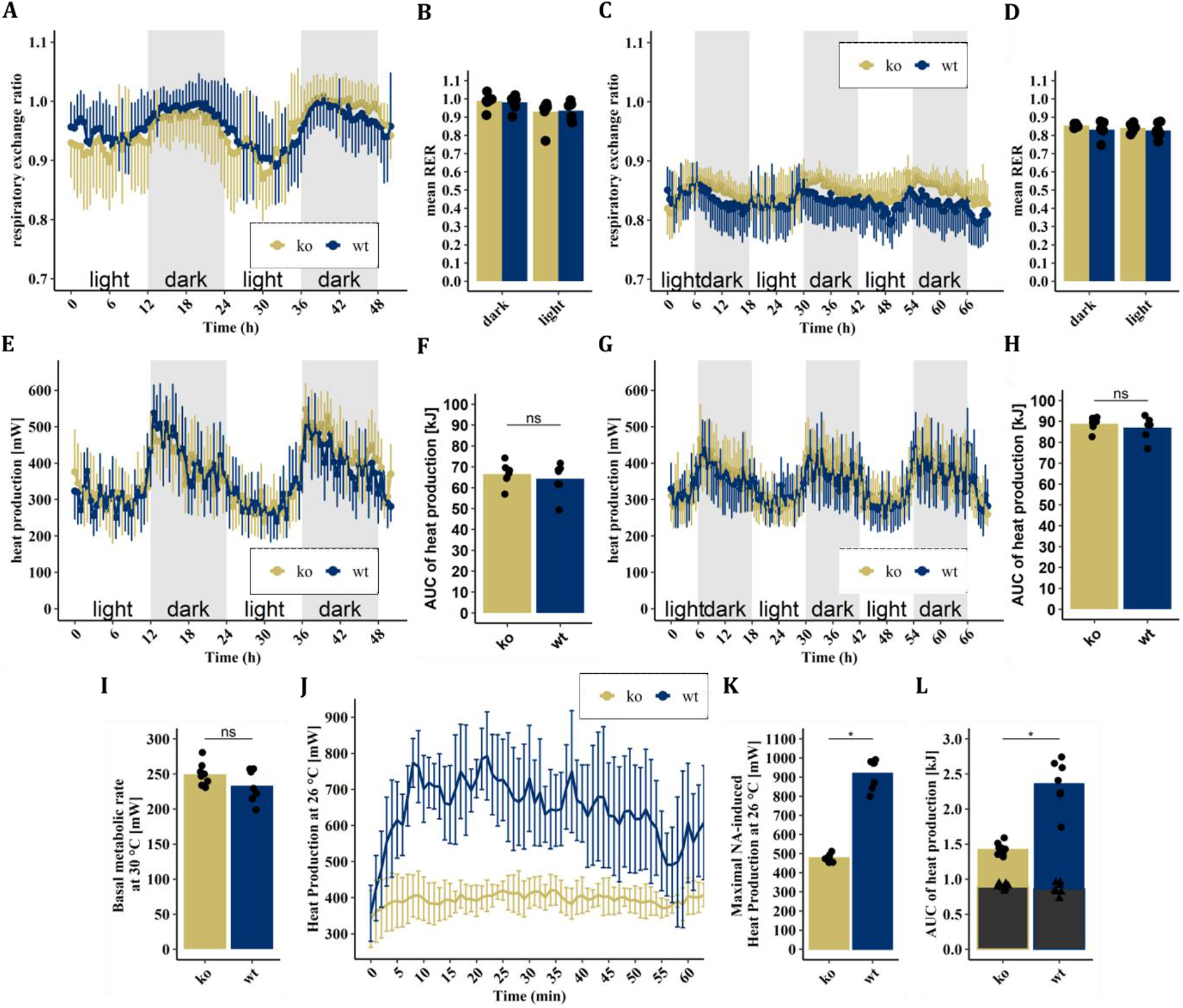
Energy expenditure at thermoneutrality is comparable between Ucp1-KO and UCP1-WT mice. **(A)** Respiratory exchange ratio of Ucp1-KO (ko, n = 7) and Ucp1-WT (wt, n = 7) mice during CD feeding. **(B)** mean respiratory exchange ratio (RER) of dark and light phases corresponding to (A). **(C)** Respiratory exchange ratio of mice during HFD feeding. **(D)** mean respiratory exchange ratio (RER) of dark and light phases corresponding to (C). **(E)** Heat production during CD feeding and **(F)** the respective area under the curve (AUC). **(G)** Heat production during HFD feeding and **(H)** the respective area under the curve (AUC). **(I)** Mean basal metabolic rate (mean of the four consecutive lowest values after at least 3h of fasting) at 30 °C. **(J)** Heat production curve of mice injected with noradrenalin at 26 °C. **(K)** Maximal heat production during the 80 minutes measurement interval shown in (J). **(L)** AUC of heat production corresponding to (J). Grey bars and triangles indicating contribution of basal metabolic rate. (F,H,I,K,L) Students t-test, ns = p > 0.5, * = p < 0.5, bars indicate group means; (A,C,E,G) data represented as means and standard deviation, averaged over a period of 30 min; (J) data represented as means and standard deviation, averaged over a period of 10 min.

The activation of BAT thermogenesis by feeding (diet-induced thermogenesis) might contribute to total energy expenditure and thus protect WT mice from diet induced obesity (von Essen et al. 2017). To scrutinize these findings, we investigated whether knockout of UCP1 affected energy expenditure in the new mouse model. As expected, metabolic rate in terms of O_2_ consumption, CO_2_ production and heat production were subject to diurnal alterations during CD and HFD feeding, increasing during the nocturnal activity phase, and decreasing during the daytime resting phase (Figure 7 E&G and Supplementary Figure 5 A-F). However, energy expenditure (area under the heat production curve) during the measurements were similar between WT and KO mice at all times (Figure 7 F&H and Supplementary Figure 5 E&F). Consequently, knockout of UCP1 did not affect energy expenditure at thermoneutral conditions.

Subsequent to the energy expenditure measurement during HFD feeding, we investigated basal metabolic rate and noradrenaline (NA) induced heat production in fasted mice. KO and WT mice hat similar basal metabolic rate (Figure 7 I) and increased metabolic rates after NA injection, similar to previous observations (Granneman et al. 2003; Meyer et al. 2010). However, WT showed a remarkably higher response upon NA injection compared to KO (Figure 7 J-L), demonstrating the capacity for UCP1 mediated thermogenesis. In addition to energy expenditure, we measured energy intake and fecal energy excretion during the calorimetry sessions. Faecal energy content was higher during high fat compared to control diet feeding, reflecting the increased energy content of the high-fat diet (not shown). However, there was no difference between genotypes in either feeding period (Figure 8 A&B). This was also true for excreted (Figure 8 C&D) and ingested (Figure 8 E&F) energy during the calorimetry sessions. Consequently, energy balance (ingested energy – excreted energy – AUC of heat production) was unaffected by the deletion of UCP1 under thermoneutral conditions (Figure 8 G&H).

**Figure 8:**
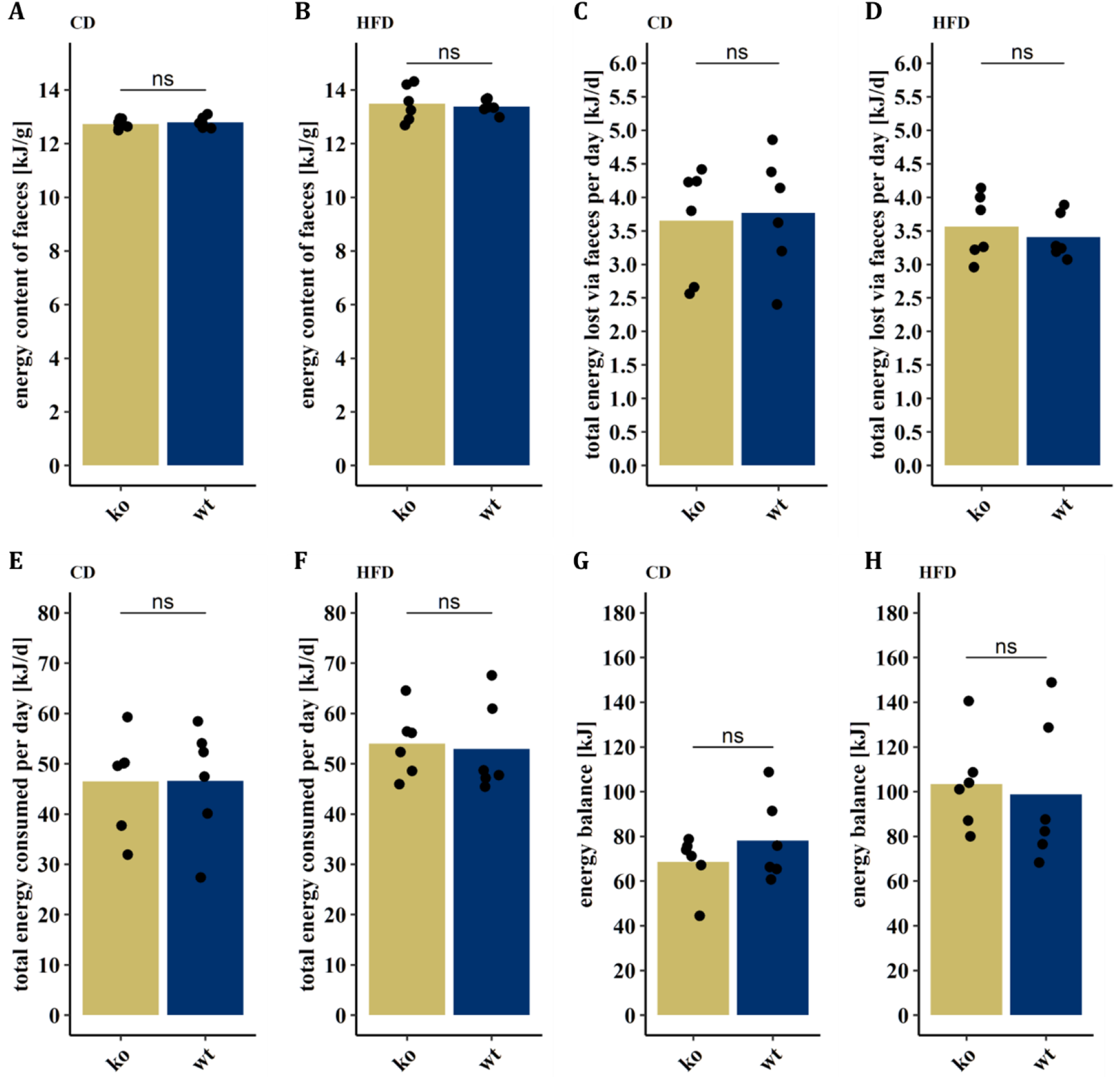
Knockout of Ucp1 does not influence energy balance at thermoneutrality. Faecal energy content of Ucp1-WT (n = 6) and Ucp1-KO (n = 6) fed **(A)** CD or **(B)** HFD. Total energy lost via faeces of mice fed **(C)** CD or **(D)** HFD. Energy consumption of mice fed **(E)** CD or **(F)** HFD. Energy balance of mice during **(G)** CD or **(H)** HFD feeding. One mouse (wt) was removed as it did not eat during the calorimetry session. Another one (ko) was removed from the analysis as the faecal samples did not combust completely. Students t-test, ns = p > 0.5, bars indicate group means.

## 4 Discussion

We characterize for the first time a novel UCP1 knockout model, as an alternative to the established and widely used UCP1-KO mouse, generated by Leslie Kozak and coworkers (Enerbäck et al. 1997). In regard of the conflicting data published so far on the DIO susceptibility of this established and widely used UCP1-KO model, there is an urgent need for new UCP1-KO models generated by cutting edge transgenic technologies to enable robust validation of metabolic functions for UCP1. Comprehensive metabolic phenotyping of the novel UCP1-KO mouse model presented in this study will help to clarify the role of UCP1 in brown and brite/beige adipose tissues for energy balance regulation, contrasting diet-and cold-induced non-shivering thermogenesis. Down this line, another UCP1-KO mouse model lacking functional UCP1 due to a SNP at nucleotide 38 of exon 5 of the UCP1 gene will also be instrumental for comparative studies (Bond and Ntambi 2018).

The aims of the study were to provide a basal characterization of the constitutive UCP1 knockout mouse and to validate this model by comparison with the results of previous studies on the established UCP1-KO mouse, especially in light of the still ongoing debate whether (Feldmann et al. 2009; von Essen et al. 2017; Rowland et al. 2016; Luijten et al. 2019; Pahlavani et al. 2019) or not (Enerbäck et al. 1997; Liu et al. 2003; Zietak and Kozak 2016; Winn et al. 2017; Maurer et al. 2020) the knockout of UCP1 renders mice more susceptible to DIO at thermoneutral conditions.

At standard housing conditions (~23°C ambient temperature) mice rely on constantly active thermogenesis to maintain normothermia. Mice lacking UCP1 recruit other thermogenic mechanisms to defend body temperature at these conditions and thus in contrast to WT mice are protected against DIO obesity (Keipert et al. 2020; T. Wang et al. 2008; Liu et al. 2003). Similarly, UCP1-KO pups showed decreased body temperature (iSST) in the present study and decreased weight after weaning, confirming the significance of UCP1 as an efficient mechanism to defend body temperature in early life. Interestingly, differences in body weight after weaning could be seen in pups of the conventional UCP1-KO mouse on a 129S1/SvImJ but not on a C57BL/6J background. Indeed, the effect of UCP1 knockout depends on the genetic background since congenic 129S1/SvImJ or C57BL/6J UCP1 knockout mice are cold sensitive, while their F1-hybrids are not (Hofmann et al. 2001).

Once the need for thermoregulatory heat production is eliminated by housing mice in their thermoneutral zone (27-30°C) the effect of UCP1 deletion becomes inconclusive. Results from several studies suggest that at thermoneutrality UCP1-KO mice are more susceptible to DIO due to the lack of diet-induced thermogenesis (von Essen et al. 2017; Feldmann et al. 2009; Rowland et al. 2016; Luijten et al. 2019), a mechanism activating BAT thermogenesis enabling rodents to increase their energy expenditure to avoid excessive weight gain caused by overfeeding (Bachman et al. 2002; Rothwell and Stock 1979). Thus, it seems plausible that mice lacking UCP1 are more susceptible to DIO at thermoneutrality (Feldmann et al. 2009; Luijten et al. 2019; von Essen et al. 2017; Rowland et al. 2016) considering UCP1 as the main contributor to BAT thermogenesis. We investigated this phenomenon by comprehensive metabolic analysis of our novel UCP1-KO model. Total energy expenditure was similar in both KO and WT mice and did not differ during the nocturnal feeding period, indicating no effect of UCP1 ablation on diet induced thermogenesis. Further, considering the similarities in body weight gain, food intake and metabolic efficiency between KO and WT animals there is no evidence for a more DIO susceptible phenotype of UCP1-KO mice. These findings are in line with various studies on the established UCP1-KO mouse (Enerbäck et al. 1997; Zietak and Kozak 2016; Winn et al. 2017; Maurer et al. 2020) and a second recently described UCP1-KO mouse (Bond and Ntambi 2018). Based on the combined evidence of different UCP1-KO models we conclude that energy balance regulation and the development of DIO are not affected by the presence of UCP1 at thermoneutrality.

For future studies, a major advantage and novelty of our mouse model compared to other available UCP1-KO models (Enerbäck et al. 1997; Bond and Ntambi 2018) is the option to induce conditional Cre-mediated deletion of Exon 2 in the UCP1 gene, using tamoxifen-or digitonin-inducible Cre-systems. Although, we described the constitutive UCP1-KO, this system enables conditional cell-type specific or age-dependent knock-out of UCP1. This is of significance to investigate the role of alternative mechanisms for non-shivering thermogenesis that might be recruited due to the lack of UCP1 in early life stages. So far, the only available inducible model was the UCP1-DTR mouse, expressing the diphtheria toxin receptor (DTR) under control of the UCP1 promoter, thus depleting UCP1 expressing cells (Rosenwald et al. 2013; Challa et al. 2020). This provided first insights about the contribution of brite adipocytes to energy expenditure (Challa et al. 2020). In contrast, our model will allow the selective ablation of UCP1 in distinct cell types while leaving these cells otherwise functional. Further research on inducible UCP1-KO mice based on our knockout strategy will help to study the recruitment of alternative thermogenic mechanism and to clarify the role of individual thermogenic adipocytes to non-shivering thermogenesis.

In summary, we provide evidence that the abundance of UCP1 does not influence energy metabolism at thermoneutrality and provide a new mouse model as foundation for a better understanding of the contribution of UCP1 in different cell types or life stages to energy metabolism.

## Acknowledgments

We thank the animal caretakers for their support with animal work. We thank Katherina Schnabl for excellent technical support with the indirect calorimetry as well as Johanna Bruder and Josef Oeckl for assisting during tissue sampling.

This work was financed by the “Nutribrite” grant (Deutsche Forschungsgemeinschaft (DFG) #KL973/13-1 & French Agence Nationale de la Recherche #ANR-15-CE14-0033), the Collaborative Research Center (DFG-CRC1371:P13), and the Else Kröner Fresenius Foundation (EKFS) to M.K.; as well as a European Commission grant EUCOMM (EU-FP6, LSHM-CT-2005-01893). Contributions by N.K. and J.K.P. are funded by the Bavarian State Ministry of Science and the Arts within the framework of the Bavarian Research Institute for Digital Transformation (bidt). J.H. was supported by a DFG grant with the project-ID: 335447727 - SFB 1328.

## Author contributions

**S.D**. conceived the study design, planned, and performed the mouse experiments, conducted molecular analyses, analyzed, and interpreted data, and wrote the manuscript.

**A.S**. assisted tissue sampling, analyzed, and interpreted microbiome data, and revised the manuscript.

**M.W**. conceived the study design, established the animal breeding.

**S.M**. contributed to data collection and revised the manuscript.

**W.W**. generated and provided the founder mice and revised the manuscript.

**S.M**. generated the founder mice and revised the manuscript.

**M.H.A**. generated the founder mice and revised the manuscript.

**R.K**. generated the founder mice and revised the manuscript.

**A.W**. performed the lipidomic analysis.

**M.F**. performed the lipidomic analysis and revised the manuscript.

**J.H**. performed the lipidomic analysis and revised the manuscript.

**N.K**. performed multi-omics analysis and revised the manuscript.

**J.P**. performed multi-omics analysis and revised the manuscript.

**M.K**. conceived the study design, contributed to the interpretation of the data, edited the manuscript, and acquired funding.

All authors approved the final version of the manuscript.

