## Supplement for "Susceptibility to diet induced obesity at thermoneutral conditions is independent of UCP1"

### Supplements

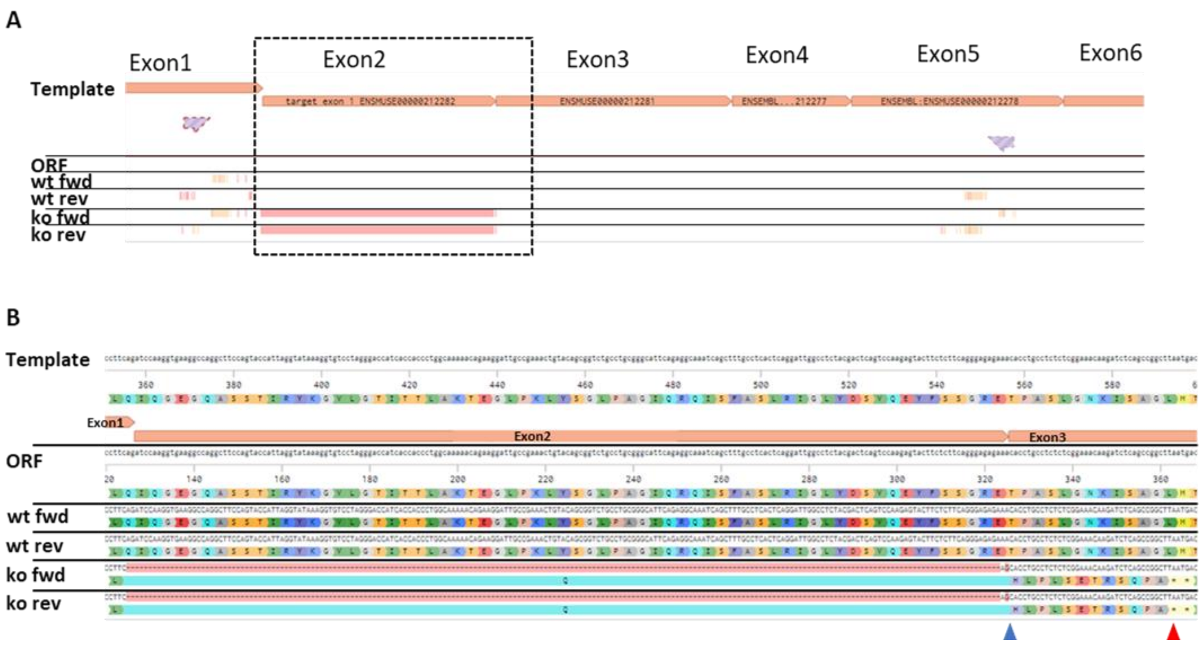


***Supplementary Figure 1: Sequencing of RT-PCR products obtained from iBAT of Ucp1-KO and Ucp1-WT mice.***

(**A**) Overview of the alignment of the Ucp1 coding sequence with open reading frame (ORF), as well as the sequencing results of Ucp1-WT (wt) and UCp1-KO (ko) for forward (fwd) and reverse (rev) primer. (**B**) Magnification of the dashed box in (A), including amino acid translation matched to the ORF. Blue and red arrowhead indicate the position of the frame shift (blue) and the premature stop codon (red).

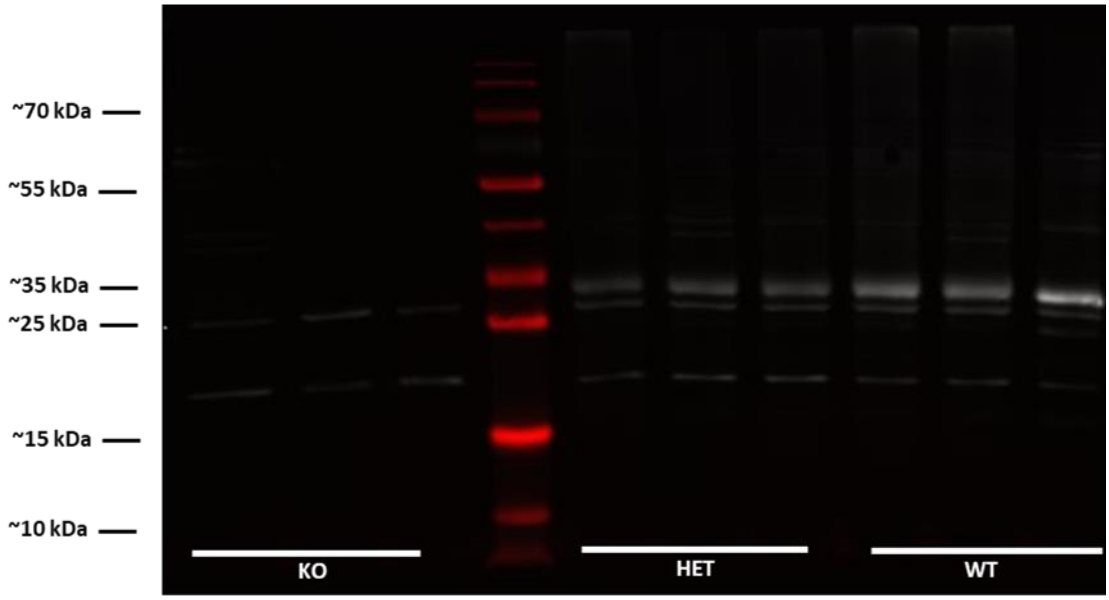


***Supplementary Figure 2: Uncropped western blot image corresponding to Figure 1D.***
800nm-Channel in white, 700nm-Channel in red.

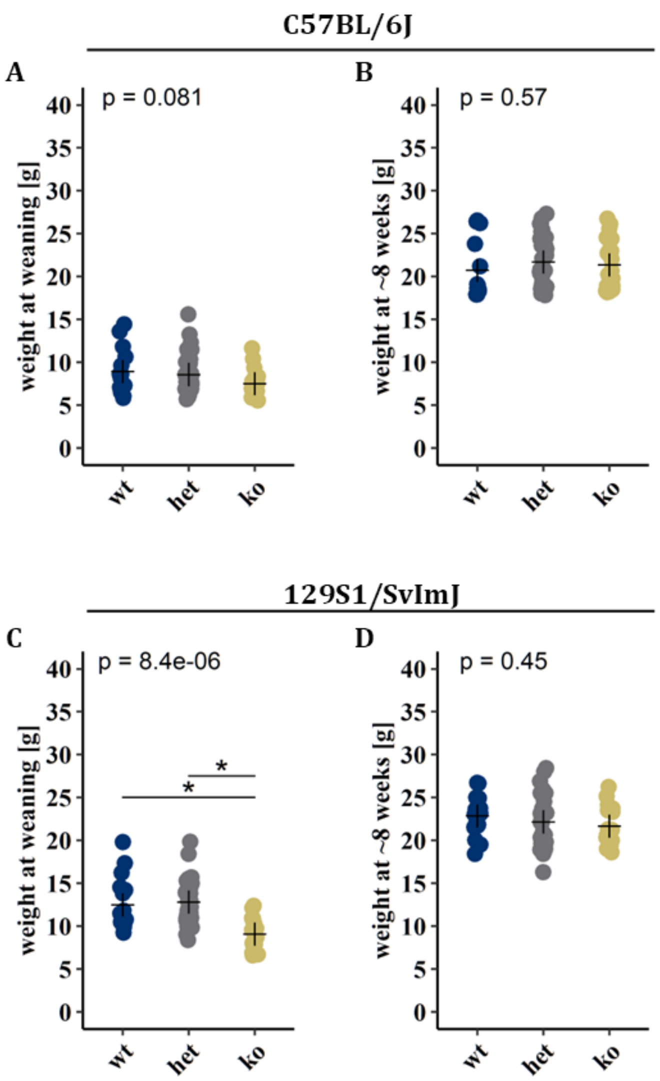


***Supplementary Figure 3: Body weights of offspring produced by HET/HET breeding of the conventional Ucp1-KO model on (A&B) C57BL/6J or (C&D) 129S1/SvImJ background.***

(A&C) at weaning ~3 weeks of age and (B&D) at the age of ~8-weeks. Crosses indicating group means.

1-Way ANOVA and t-test with bonferroni adjusted p-value, * = p-value < 0.05

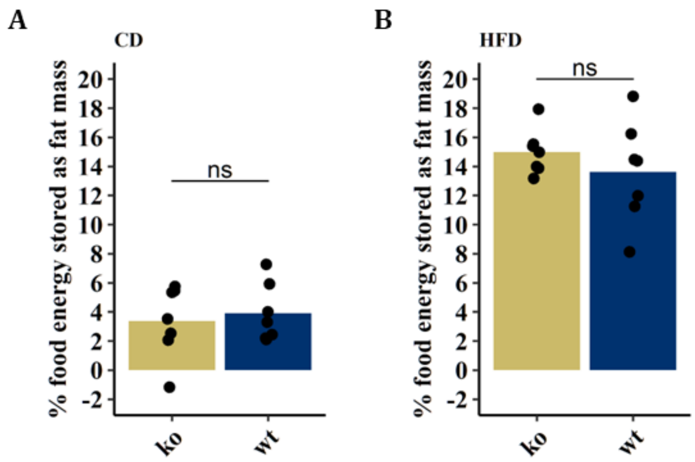


***Supplementary Figure 4: Metabolic efficiency expressed as percent of the total food energy consumed stored as fat mass during (A) CD and (B) HFD feeding.***

Sudents t-test between Ucp1-WT (wt n = 7) and Ucp1-KO (ko n = 7), ns = $p > 0.5$, bars indicate group means. Metabolic efficacy expressed as percentage of food energy stored as fat mass was calculated according to (Von Essen 2017). Briefly total fat mass gain (g) during each period was multiplied by 37.4 kJ/g and divided by food intake (kJ).

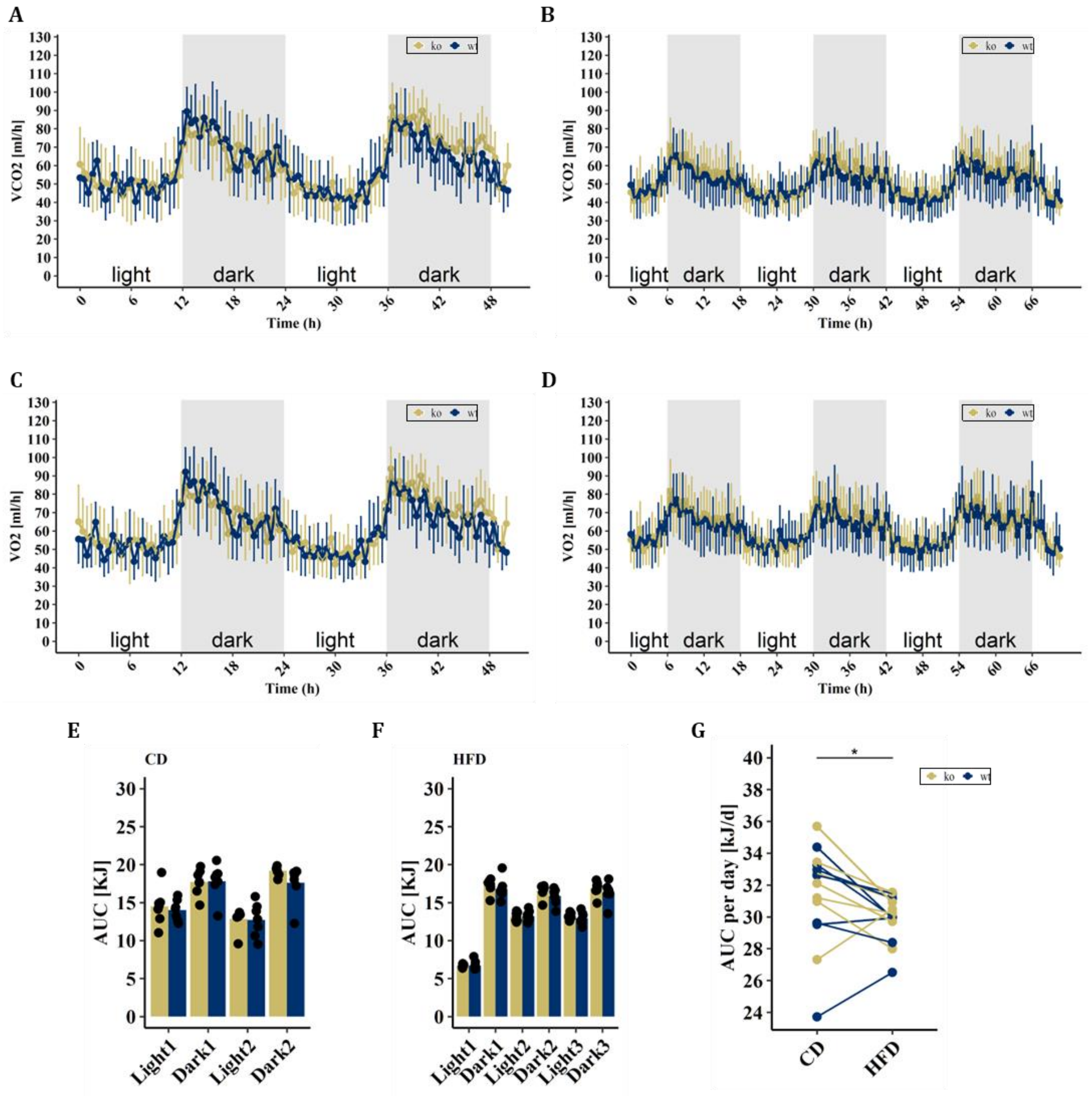


***Supplementary Figure 5: (A&B) CO2 production and (C&D) VO2 consumption during (A,C) CD and (B,D) HFD feeding.***

Area under the curve (AUC) of heat production during the different light phases corresponding to (E) Figure 4E and (F) Figure 4G. (G) Area under the curve (AUC) of heat production corresponding to Figure 4E (CD) and Figure 4G (HFD) per day. (A-D) Data represented as means and standard deviation, averaged over a period of 30 min. (E,F) Bars indicate group means. (G) Paired students t-test, * = p-value < 0.05
